# Cardiomyocyte-Specific Deletion of GCN5L1 in Mice Limits Ex Vivo Cardiac Functional Decline in Response to a High Fat Diet

**DOI:** 10.1101/805283

**Authors:** Dharendra Thapa, Janet R Manning, Michael W. Stoner, Manling Zhang, Bingxian Xie, Michael N. Sack, Iain Scott

**Affiliations:** Vascular Medicine Institute, Department of Medicine, University of Pittsburgh, Pittsburgh, PA 15261; Center for Metabolism and Department of Medicine, University of Pittsburgh, Pittsburgh, PA 15261; Division of Cardiology, Department of Medicine, University of Pittsburgh, Pittsburgh, PA 15261; Laboratory of Mitochondrial Biology, Division of Intramural Research, National Heart, Lung and Blood Institute, Bethesda, MD 20892

**Author notes:** Corresponding Author, Address for Correspondence: Iain Scott, PhD, Division of Cardiology, University of Pittsburgh, BST E1256, 200 Lothrop Street, Pittsburgh, PA, 15261.

**Keywords:** GCN5L1, Mitochondrial Acetylation, Pyruvate Utilization, Reactive Oxygen Species, Lipid Peroxidation

## Abstract

Reduced substrate flexibility, and a shift towards increased cardiac fatty acid utilization, is a key feature of obesity and diabetes. We previously reported that increased acetylation of several mitochondrial FAO enzymes, regulated in part via increased abundance of the mitochondrial acetyltransferase GCN5L1, correlated with increased FAO enzyme activity in the heart. The focus of the current study was to investigate whether decreased acetylation, via cardiomyocyte-specific deletion of GCN5L1 (GCN5L1 cKO), regulates cardiac energy metabolism following exposure to a high fat diet (HFD). Excess dietary fat led to similar cardiac hypertrophy in wildtype and GCN5L1 cKO mice. We show that acetylation of pyruvate dehydrogenase (PDH) was significantly reduced in HFD GCN5L1 cKO hearts, which correlated with its increased enzymatic activity relative to HFD wildtype controls. The acetylation of both electron transport chain Complex I protein NDUFB8, and manganese superoxide dismutase 2 (SOD2), was significantly reduced by GCN5L1 deletion in HFD animals, resulting in decreased lipid peroxidation. Finally, we show that in contrast to wildtype mice, GCN5L1 cKO hearts maintain *ex vivo* contractility and workload in response to a HFD. In summary, we show that GCN5L1 deletion limits cardiac functional decline observed in HFD mice, by increasing fuel substrate flexibility and limiting reactive oxygen species damage.

## INTRODUCTION

A healthy heart uses several fuel substrates to power contractile function [1]. Fatty acid oxidation (FAO) accounts for ~70% of total ATP production [2–4], with the remainder produced from other sources such as glucose, lactose and ketones [5]. The ability of the heart to use various fuel substrates is lost in disease states such as diabetes, which features a shift towards increased fatty acid utilization and decreased glucose oxidation [6]. While controversy exists about the possible protective role of cardiomyocyte insulin resistance in the failing heart [7], the over reliance on fatty acids during diabetes results in increased myocardial lipid accumulation, cardiac dysfunction, and ultimately decreased cardiac efficiency [8–11]. In this context, identifying mechanisms that regulate substrate utilization and cardiac energy metabolism in obesity and diabetes is of prime importance.

Growing evidence pinpoints to a crucial role played by lysine acetylation, a reversible post-translational modification, in the regulation of fuel utilization and mitochondrial bioenergetics. A study by Pougovkina et al. has shown that circulating free fatty acids, highly prevalent in obesity and diabetes, are the main source of the acetyl groups used for lysine acetylation [12]. Furthermore, we and several others have shown that mitochondrial protein acetylation is significantly increased in various tissues during obesity [13–16]. Specifically in the heart, we have shown that an increase in acetylation results in increased activity of several FAO enzymes, which is regulated in part by the mitochondrial acetyltransferase protein GCN5L1 [16]. These findings place nutritional inputs, lysine acetylation, and GCN5L1 at a key regulatory node in cardiac fuel metabolism, and hence may play a central role in the development of metabolic dysfunction.

GCN5L1 is a mitochondrial matrix protein that has been shown to counter the activity of the mitochondrial deacetylase enzyme SIRT3. Specifically, it has been shown to regulate the acetylation of several mitochondrial FAO, glucose oxidation, and electron transport chain (ETC) proteins [15–17]. Loss of GCN5L1 has been shown to limit mitochondrial respiratory capacity [18], while increasing glucose utilization and lowering fatty acid use [16, 19]. While the effect of GCN5L1 depletion in the control of fuel substrate metabolism has been studied *in vitro*, it is not known whether this regulation is operable *in vivo*, or whether changes in nutrient conditions affect functional outputs.

In the present study, we examined the effect of a long-term high fat diet (HFD) in wildtype (WT) and cardiac-specific GCN5L1 knockout (cKO) mice. Despite similar levels of cardiac hypertrophy, GCN5L1 cKO mice maintain cardiac workload *ex vivo* after HFD exposure, while WT mice decreased output by ~50% under the same conditions. We show that GCN5L1 cKO mice display elevated pyruvate dehydrogenase (PDH) activity under HFD conditions relative to WT mice, which correlates with decreased enzyme acetylation. GCN5L1 expression regulates the acetylation of the ETC Complex I protein NDUFB8, as well as mitochondrial superoxide dismutase 2 protein (SOD2), resulting in decreased reactive oxygen species (ROS) damage in GCN5L1 cKO mice following HFD exposure. Taken together, these data indicate that acetylation of cardiac mitochondrial proteins by GCN5L1 under HFD conditions results in cardiac dysfunction via reduced fuel substrate flexibility and increased ROS production.

## MATERIALS AND METHODS

### Animal Care and Use

Animals were housed in the University of Pittsburgh animal facility under standard conditions with *ad libitum* access to water and food, and maintained on a constant 12h light/12h dark cycle. Male GCN5L1 WT and cKO animals were fed either a standard low fat diet (LFD; 70% carbohydrate, 20% protein, 10% fat; Research Diets D12450B), or a high fat diet (HFD; 20% carbohydrate, 20% protein, 60% fat; Research Diets D12492), for 24 weeks. At the end of 24 weeks, animals were euthanized and heart tissues excised for analysis. Experiments were conducted in compliance with National Institutes of Health guidelines, and followed procedures approved by the University of Pittsburgh Institutional Animal Care and Use Committee.

### Transgenic mouse generation

Cardiomyocyte-specific inducible GCN5L1 knockout mice used in the studies were generated as previously reported [20]. Briefly, mice were generated on a C57BL/6J background by crossing αMHC-Cre (Jax B6.FVB(129)-Tg(Myh6-cre/Esr1*)1JMK/J) mice (i.e. αMHC-MerCreMer) to mice with LoxP sites introduced around exon 3 of Bloc1s1 (GCN5L1^FL)^. Cardiomyocyte-specific GCN5L1 deletion was induced via tamoxifen injection (single 40 mg/kg IP injection) and confirmed by qPCR and immunoblot.

### Protein Isolation

For whole heart protein lysate, tissues were minced and lysed in CHAPS buffer on ice for ~2 hours. Homogenates were spun at 10,000 *g*, and supernatants collected for western blotting or co-immunoprecipitation experiments. For activity assays, muscle homogenates were prepared in a modified Chappell-Perry Medium A buffer (120 mM KCl, 20mM Hepes, 5 mM MgCl_2_, 1 mM EGTA, ~pH 7.2) supplemented with 5 mg/ml fat-free bovine serum albumin. After centrifugation at 10,000 *g*, supernatants were used to assay enzyme activity. Mitochondria isolation was performed using the Qproteome Mitochondria isolation kit (Qiagen). Briefly, heart tissues were cut and homogenized in the provided lysis buffer. After separating the cytoplasmic fraction by centrifuging at 1000 *g* at 4 °C for 10 minutes, the remaining pellet was suspended in disruption buffer and centrifuged at 1000 *g* at 4 °C for 10 minutes. The supernatant was retained and centrifuged at 6000 *g* at 4 °C for 10 minutes to obtain the mitochondrial pellet. Mitochondria were reconstituted in the appropriate assay buffer depending upon the experiments performed.

### Western Blotting

Protein lysates were prepared in LDS sample buffer, separated using Bolt SDS/PAGE 4-12% or 12% Bis-Tris gels, and transferred to nitrocellulose membranes (all Life Technologies). Protein expression was analyzed using the following primary antibodies: rabbit acetyl-lysine (Ac-K), rabbit glutamate dehydrogenase (GDH), and rabbit PDH from Cell Signaling Technologies; rabbit acetylated SOD2 (K122), rabbit SOD2, and rabbit OXPHOS cocktail (to analyze NDUFB8, SDHB and UQCR2 protein content) from Abcam. Fluorescent anti-mouse or anti-rabbit secondary antibodies (red, 700 nm; green, 800 nm) from LiCor were used to detect expression levels. Protein densitometry was measured using Image J software (National Institutes of Health, Bethesda, MD).

### Co-Immunoprecipitation

For co-immunoprecipitation experiments, protein lysates were harvested in CHAPS buffer, and equal amounts of protein were incubated overnight at 4 °C with rabbit acetyl-lysine antibody (AcK; Cell Signaling). Immunocaptured proteins were isolated using Protein-G agarose beads (Cell Signaling Technology), washed multiple times with CHAPS buffer, and then eluted in LDS sample buffer at 95 °C. Samples were separated on 12% Bis-Tris Bolt gels and probed with appropriate antibodies. Protein densitometry was measured using Image J software (National Institutes of Health, Bethesda, MD).

### Isolated Working Heart

Cardiac contraction and workload were calculated using the isolated working heart system as previously described [20]. Hearts from anesthetized mice were rapidly excised and cannulated via the aorta in warm oxygenated Krebs-Henseleit buffer (118 mM NaCl, 25 mM NaHCO_3_, 0.5 mM Na-EDTA [disodium salt dihydrate], 5 mM KCl, 1.2 mM KH_2_PO_4_, 1.2 mM MgSO_4_, 2.5 mM CaCl_2_, 11 mM glucose). Retrograde (i.e. Langendorff) perfusion was initiated to blanch the heart, maintained at a constant aortic pressure of 50 mmHg with a peristaltic pump through a Starling resistor. A small incision was next made in the pulmonary artery to allow perfusate to drain, and the heart was paced at a rate slightly higher than endogenous (~360-500 bpm). The left atrium was then cannulated via the pulmonary vein, and anterograde perfusion was initiated with a constant atrial pressure of 11 mmHg against an aortic workload of 50 mmHg. Left ventricle pressure was measured via Mikro-tip pressure catheter (Millar) carefully inserted into the LV through the aorta. The work-performing heart was permitted to equilibrate for 30 minutes to establish baseline functional parameters.

*Ex vivo* cardiac workload was calculated as the difference between recorded atrial and aortic pressures, multiplied by the cardiac output flow parameter. Workload was then normalized for each heart by dry heart weight (determined by measuring the ratio of a small section of air-dried heart tissue to its wet weight, and multiplying the entire heart wet weight by that ratio), and expressed as mmHg/mL/min/g. The difference in workload between LFD and HFD was determined as the percent normalized workload of the HFD relative to LFD for each genotype.

### Biochemical Assays

PDH activity and lipid peroxidation were measured using commercial kits (Sigma-Aldrich) according to manufacturer’s instructions.

### Statistics

Graphpad Prism software was used to perform statistical analyses. Means ± SE were calculated for all data sets. Data were analyzed using two-way ANOVA with Sidak post-hoc testing to determine differences between genotypes and feeding groups. Data were analyzed with two-tailed Student’s T-Tests to determine differences between single variable groups. *P* < 0.05 was considered statistically significant.

## RESULTS

### Loss of GCN5L1 decreases mitochondrial protein acetylation under high fat diet conditions

We first examined the response of WT and GCN5L1 cKO mice to varying nutrient conditions, and placed groups of animals on a LFD or HFD for 24 weeks. Mice exposed to a HFD displayed a significant increase in body weight and heart weight in both WT and GCN5L1 cKO animals (Fig 1A-B). We previously reported that the increased acetylation and activity of several mitochondrial FAO enzymes following HFD feeding is regulated in part by GCN5L1, using *in vitro* cell models [16]. To further investigate this role of GCN5L1 *in vivo*, we first examined the total level of mitochondrial protein acetylation in WT and GCN5L1 cKO animals under LFD and HFD. While still significantly elevated relative to LFD mice, there was a significant ~67% decrease in mitochondrial acetylation in GCN5L1 cKO HFD mice relative to their WT HFD controls (Fig 1C-D). We therefore conclude that while loss of GCN5L1 does not affect major phenotypic responses to excess nutrition, its deletion limits the accumulation of hyperacetylated mitochondrial proteins.

**Figure 1:**
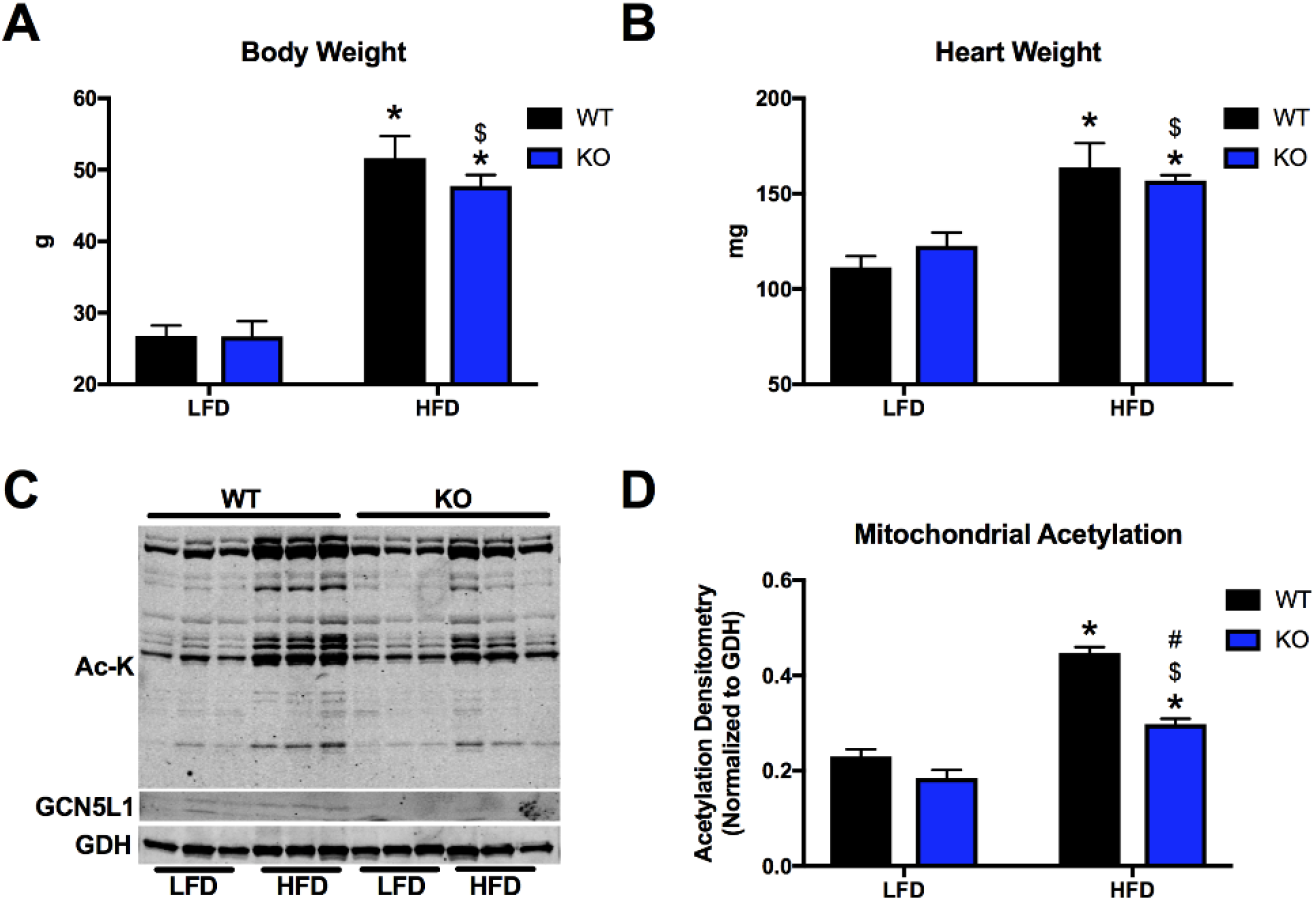
Characterization of mitochondrial acetylation in GCN5L1 WT and cKO animals. (A-B) Significant increases in body weight and wet heart weight were observed in both WT and cKO HFD animals, with no significant difference detected between genotypes. (C-D) Overall cardiac mitochondrial protein acetylation was greatly increased in HFD WT animals, which was significantly attenuated in HFD cKO mice. Values are expressed as means ± SEM, n = 5-6, * *P* < 0.05 vs WT LFD, ^$^ *P* < 0.05 vs cKO LFD, ^#^ *P* < 0.05 vs WT HFD.

### GCN5L1 regulates the acetylation and enzymatic activity of PDH

After examining the role of GCN5L1 in the regulation of global mitochondrial protein acetylation, we next examined whether its deletion had any effect on the acetylation of key metabolic regulatory proteins. Acetylation of pyruvate dehydrogenase (PDH) was significantly decreased in GCN5L1 cKO HFD animals when compared to their WT HFD controls (Fig 2A-B). Acetylation of PDH has been shown to inhibit its activity in the heart [21]. While it did not reach statistical significance (*P* = 0.12), we observed that PDH activity in GCN5L1 cKO HFD animals was 88% higher than in WT HFD mice (Fig 2C). Furthermore, linear regression analyses between PDH acetylation status and enzyme activity showed that there was a significant negative correlation between PDH activity and acetylation (*r^2^* = 0.3352, *P* = 0.03; Fig 2D). We therefore conclude that loss of GCN5L1 in the heart may allow mitochondrial pyruvate utilization to continue under HFD conditions by maintaining PDH activity at a similar rate to that found in LFD animals.

**Figure 2:**
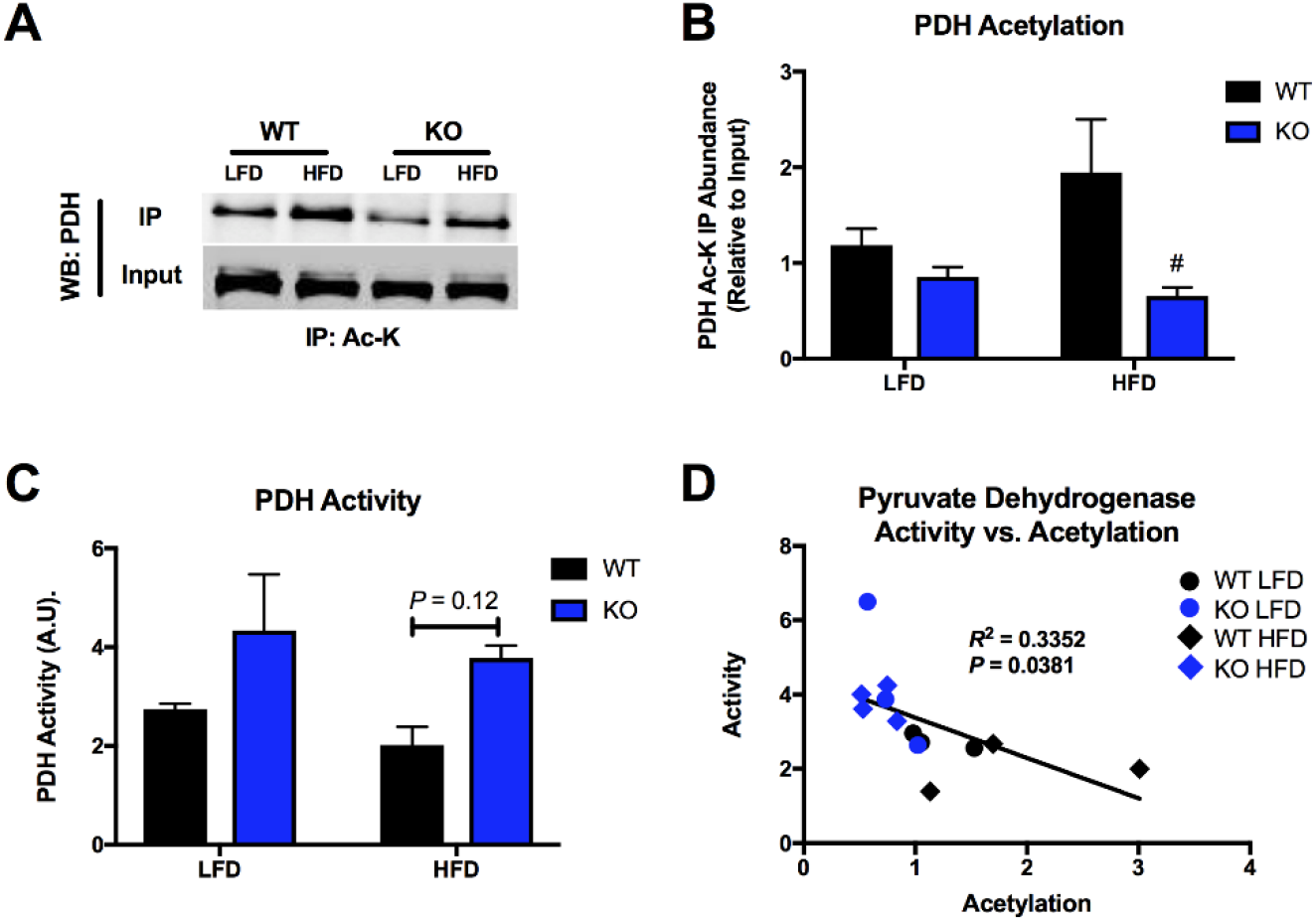
Impact of GCN5L1 deletion on the acetylation and activity of PDH. (A-B) Increased PDH acetylation levels observed in WT HFD animals were absent in cKO HFD mice. (C) Under HFD conditions, PDH activity was 88% higher in cKO animals relative to WT mice, although this did not reach statistical significance (*P* = 0.12). (D) Linear regression analyses showed a significant negative correlation between PDH acetylation status and enzymatic activity. Values are expressed as means ± SEM, n = 3-5, ^#^ *P* < 0.05 vs WT HFD.

### Role of GCN5L1 in electron transport chain (ETC) complex acetylation and redox status

We next examined the role of GCN5L1 in the acetylation of ETC complex proteins. Immunoprecipitation with an acetylated lysine antibody showed a significant increase in the acetylation of Complex I subunit NADH dehydrogenase [ubiquinone] 1 beta subcomplex subunit 8 (NDUFB8) in GCN5L1 WT HFD animals, which was completely attenuated in GCN5L1 cKO mice (Fig 3A). No significant changes were observed in acetylation of Complex II and III subunits SDHB and UQCR2, respectively (Fig 3B-C).

**Figure 3:**
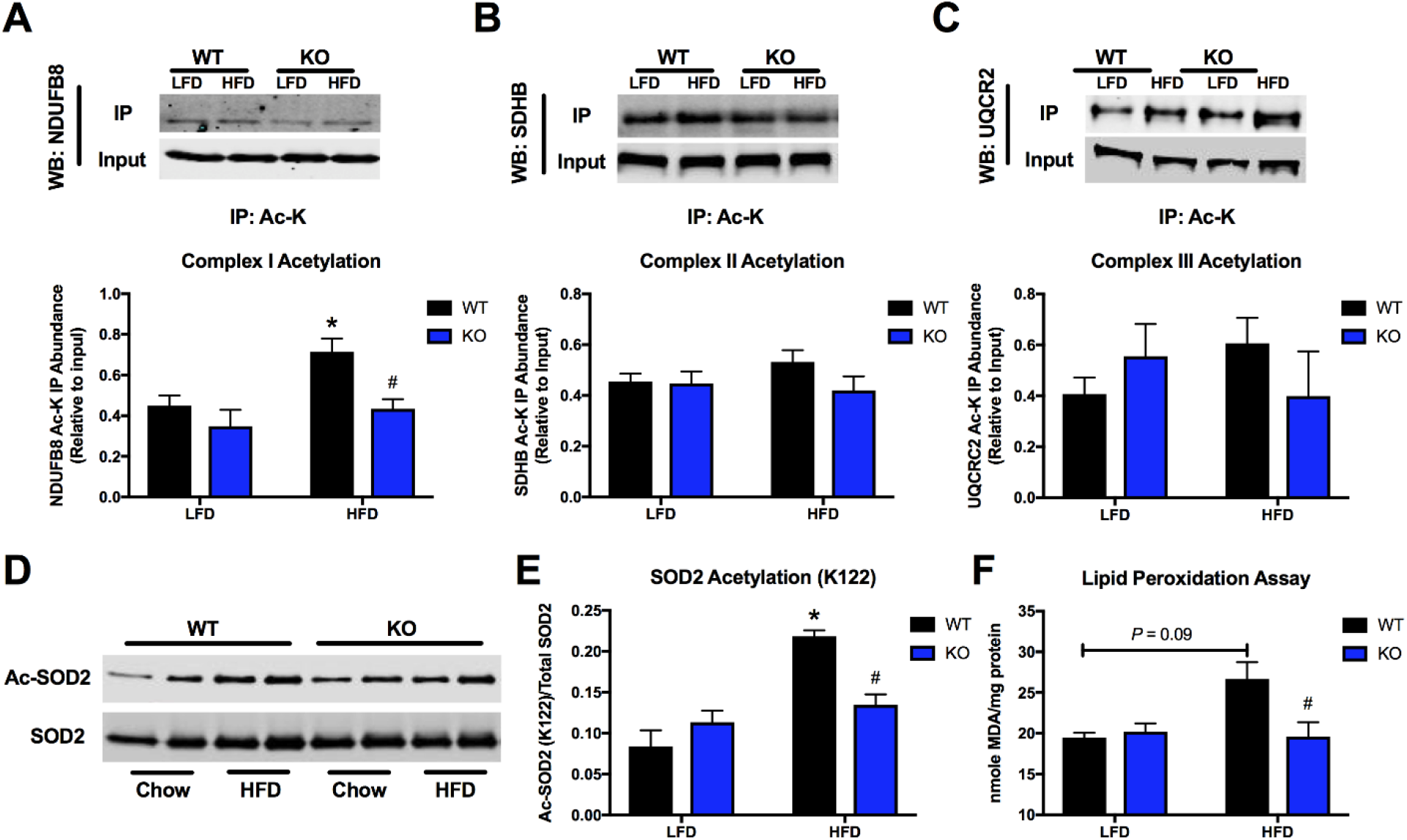
Acetylation status of electron transport chain and ROS regulatory proteins. (A) Acetylated lysine immunoprecipitation showed an increase in the acetylation of NDUFB8 (Complex I) in WT HFD animals relative to cKO mice under the same conditions. (B-C) There were no changes observed in the acetylation of SDHB (Complex II) and UQCR2 (Complex III). (D-E) Significant increases in acetylated SOD2 (K122) observed in WT HFD animals were absent in cKO HFD mouse hearts. (F) Significant increases in lipid peroxidation observed with WT HFD mice were absent in cKO HFD mice. Values are expressed as means ± SEM, n = 5-6, * *P* < 0.05 vs WT LFD, ^#^ *P* < 0.05 vs WT HFD.

ETC Complex I is a major source of reactive oxygen species production [22]. We next investigated whether changes in the acetylation of Complex I proteins impacted the redox milieu in GCN5L1 cKO mice. Deacetylation of mitochondrial superoxide dismutase 2 (SOD2) at lysine 122 (K122) has been shown to increase its activity [23]. With HFD, we observed a significant increase in the acetylation of SOD2 at K122 in WT mice, which was significantly reduced in GCN5L1 cKO animals (Fig 3D-E). Finally, we observed that an increase in lipid peroxidation in WT HFD animals was significantly attenuated in GCN5L1 cKO mice (Fig 3F). We therefore conclude that acetylation of SOD2 is in part regulated by GCN5L1, and that loss of this modification in GCN5L1 cKO animals results in decreased lipid peroxidation in response to HFD exposure.

### Loss of GCN5L1 maintains ex vivo cardiac workload under high fat diet conditions

We hypothesized that by maintaining glucose oxidation capacity and reducing ROS damage, GCN5L1 cKO hearts may be better able to maintain cardiac work under HFD conditions. Measurement of cardiac function *ex vivo* using an isolated working heart system showed that there were no significant changes in cardiac contractility or relaxation (Fig 4A-B) between GCN5L1 WT and cKO animals under LFD or HFD conditions. Under LFD conditions, GCN5L1 cKO mice displayed a marginally lower normalized *ex vivo* cardiac workload than WT mice when perfused with glucose as a substrate, however this did not reach statistical significance (*P* = 0.25; Fig 4C). Under HFD conditions, WT mice displayed a significantly reduced *ex vivo* cardiac workload compared to WT LFD controls, while no change was observed in cKO mice between low fat and high fat conditions (Fig 4C). We therefore conclude that loss of mitochondrial protein acetylation in GCN5L1 cKO hearts allows these mice to maintain cardiac output under excess nutrient conditions that negatively impact WT mice.

**Figure 4:**
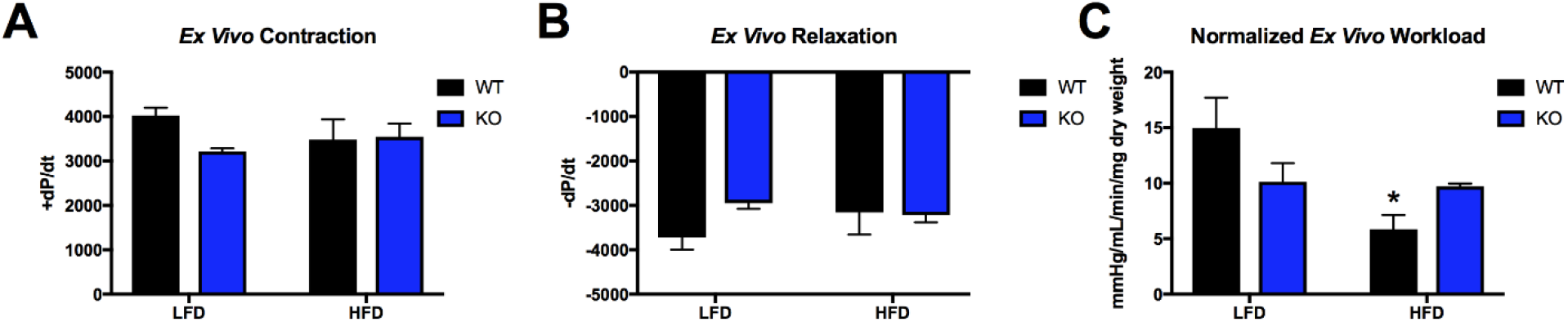
GCN5L1 cKO mice maintain cardiac function *ex vivo* under HFD conditions. (A-C) No significant changes were observed in contractility or relaxation between WT and cKO mice perfused with glucose under isolated working heart conditions. However, there was a significant decrease in normalized *ex vivo* workload in WT HFD animals relative to WT LFD mice, which was absent in cKO mice. Values are expressed as means ± SEM, n = 4-8, * *P* < 0.05 vs WT LFD.

## DISCUSSION

Using a HFD feeding model combined with cardiac-specific deletion of GCN5L1, we demonstrate for the first time *in vivo* that GCN5L1 mediates the acetylation and enzymatic activity of PDH in response to nutrient excess. Furthermore, we show that increased acetylation of the Complex I protein NDUFB8, and the superoxide dismutase enzyme SOD2, is absent in GCN5L1 cKO animals, which correlates with decreased lipid peroxidation under HFD conditions. Finally, we show that a significant decrease in *ex vivo* cardiac workload found in WT animals under HFD conditions is not observed in GCN5L1 cKO mice. In summary, we conclude that GCN5L1 deletion limits cardiac functional decline observed in mice under conditions of nutrient excess.

We have previously shown that prolonged high fat feeding leads to the increased acetylation of mitochondrial proteins, which was in part regulated by GCN5L1 when tested under *in vitro* cell culture conditions [16]. To further substantiate the role played by GCN5L1 in regulating the mitochondrial acetylome following exposure to a HFD, we utilized a newly described cardiac-specific GCN5L1 cKO animal model [20]. As anticipated, we observed a significant decrease in mitochondrial protein acetylation in GCN5L1 cKO animals under nutrient excess conditions. A number of studies have extensively reported the role of SIRT3-mediated mitochondrial protein deacetylation [14, 24, 25], however the role of GCN5L1 in this context is much less known. Upon further investigation of specific proteins targeted by GCN5L1, we found that increased acetylation of PDH is greatly attenuated by GCN5L1 deletion under HFD conditions.

We have previously shown that genetic depletion of GCN5L1 in rat H9c2 and human AC16 cardiac cells promotes an increase in glucose utilization for bioenergetic use [18, 19]. This increase did not affect basal respiration in either cell type relative to control cells, however maximal respiration capacity in response to chemical uncoupling was reduced in GCN5L1 knockdown cells under high glucose conditions [18, 19]. In the present study, we show that PDH activity (a major regulator of mitochondrial glucose utilization) is maintained at a higher level in GCN5L1 cKO cardiac tissue than WT mice under both LFD and HFD conditions (Fig 2). This suggests that *in vivo* deletion of GCN5L1 in the heart will result in higher glucose utilization capacity, matching what we observed *in vitro*. However, a key difference based may be that GCN5L1 cKO mice are able to use glucose to drive oxidative phosphorylation, while GCN5L1 knockdown in cell culture drives metabolism instead towards an oxidative glycolytic phenotype [19]. Studies are currently underway to measure cardiac fuel substrate utilization *in vivo*, which will determine which glucose utilization pathway predominates in the GCN5L1-depleted heart.

It was previously reported that ETC Complex I is one of the major sites of ROS production in the mitochondria [26–28]. Complex I deficiency has also been shown to increase mitochondrial protein acetylation and accelerate heart failure, via SIRT3-mediated hyperacetylation [29]. Furthermore, in a streptozotocin-induced type 1 diabetic model, enhanced mitochondrial protein lysine acetylation has been reported as a common consequence of increased FAO and Complex I defects [30], which could ultimately be responsible for metabolic inflexibility of the diabetic heart. These authors reported no change in the mitochondrial deacetylase SIRT3, while the role of GCN5L1 was not examined [30]. We report for the first time that mitochondrial GCN5L1 partly regulates the acetylation of the Complex I protein subunit NDUFB8, which may contribute to complex activity defects under HFD conditions.

Superoxide dismutase 2 (SOD2) is the primary mitochondrial ROS scavenger that converts superoxide (a by-product of Complex I activity) to hydrogen peroxide, which is ultimately converted to water by catalase and other peroxidases [31]. It has been reported that SIRT3-mediated deacetylation of SOD2 at K122 results in its increased enzymatic activity [23]. As such, we examined if the mitochondrial acetyltransferase activity of GCN5L1 regulated this acetylation site in our study, and found that deletion of GCN5L1 resulted in significantly decreased acetylation of SOD2 at K122 in cKO HFD mice (Fig 3D-E). This may contribute to the significantly decreased lipid peroxidation observed in GCN5L1 cKO HFD hearts (Fig 3F). These new data stand in partial contrast to previous findings, which showed that GCN5L1 cKO mouse hearts displayed elevated levels of protein carbonylation (a marker of ROS damage) in an *ex vivo* model of cardiac ischemia [20]. This ROS damage led to a decline in contractile function under ischemic conditions, which could be rescued by supplying the *ex vivo* heart with the ROS scavenger n-acetyl cysteine [20]. Further studies will be required to understand the mechanistic differences underlying the response of GCN5L1-depleted hearts to ROS from different sources. Along these lines, work is currently underway to elucidate the functional outcome of K122 acetylation on SOD2 activity in response to acute (ischemia-reperfusion) vs. chronic (HFD) derived ROS in the heart. It is expected that delineation of the response to ROS from these two etiologies may explain the partial differences observed between our ischemic and HFD studies.

Based upon our findings, we conclude that cardiomyocyte-specific deletion of GCN5L1 impacts the acetylation and activity of PDH. Furthermore, we report a novel role for GCN5L1 in regulating SOD2 and NDUFB8 acetylation. These findings further substantiate our growing appreciation of GCN5L1 as a regulator of mitochondrial protein activity, and its subsequent role in controlling cardiac work. Our data suggest that by regulating the ability of the heart to use different fuel substrates, GCN5L1 may have the capacity to be a significant modulator of cardiac function in health and disease states.

## REFERENCES

1. Lopaschuk, G.D., et al., Myocardial fatty acid metabolism in health and disease. Physiol Rev, 2010. 90(1): p. 207–58.

2. Bing, R.J., et al., Metabolism of the human heart. II. Studies on fat, ketone and amino acid metabolism. Am J Med, 1954. 16(4): p. 504–15.

3. Lopaschuk, G.D., et al., Regulation of fatty acid oxidation in the mammalian heart in health and disease. Biochim Biophys Acta, 1994. 1213(3): p. 263–76.

4. Neely, J.R. and H.E. Morgan, Relationship between Carbohydrate and Lipid-Metabolism and Energy-Balance of Heart-Muscle. Annual Review of Physiology, 1974. 36: p. 413–459.

5. Doenst, T., T.D. Nguyen, and E.D. Abel, Cardiac metabolism in heart failure: implications beyond ATP production. Circ Res, 2013. 113(6): p. 709–24.

6. Lopaschuk, G.D., C.D. Folmes, and W.C. Stanley, Cardiac energy metabolism in obesity. Circ Res, 2007. 101(4): p. 335–47.

7. Taegtmeyer, H., et al., Insulin resistance protects the heart from fuel overload in dysregulated metabolic states. Am J Physiol Heart Circ Physiol, 2013. 305(12): p. H1693-7.

8. Carley, A.N., et al., Mechanisms responsible for enhanced fatty acid utilization by perfused hearts from type 2 diabetic db/db mice. Arch Physiol Biochem, 2007. 113(2): p. 65–75.

9. Fillmore, N., J. Mori, and G.D. Lopaschuk, Mitochondrial fatty acid oxidation alterations in heart failure, ischaemic heart disease and diabetic cardiomyopathy. Br J Pharmacol, 2014. 171(8): p. 2080–90.

10. Kalaivanisailaja, J., V. Manju, and N. Nalini, Lipid profile in mice fed a high-fat diet after exogenous leptin administration. Pol J Pharmacol, 2003. 55(5): p. 763–9.

11. Young, M.E., et al., Impaired long-chain fatty acid oxidation and contractile dysfunction in the obese Zucker rat heart. Diabetes, 2002. 51(8): p. 2587–95.

12. Pougovkina, O., et al., Mitochondrial protein acetylation is driven by acetyl-CoA from fatty acid oxidation. Hum Mol Genet, 2014. 23(13): p. 3513–22.

13. Alrob, O.A., et al., Obesity-induced lysine acetylation increases cardiac fatty acid oxidation and impairs insulin signalling. Cardiovasc Res, 2014. 103(4): p. 485–97.

14. Hirschey, M.D., et al., SIRT3 regulates mitochondrial fatty-acid oxidation by reversible enzyme deacetylation. Nature, 2010. 464(7285): p. 121–5.

15. Thapa, D., et al., The protein acetylase GCN5L1 modulates hepatic fatty acid oxidation activity via acetylation of the mitochondrial beta-oxidation enzyme HADHA. J Biol Chem, 2018. 293(46): p. 17676–17684.

16. Thapa, D., et al., Acetylation of mitochondrial proteins by GCN5L1 promotes enhanced fatty acid oxidation in the heart. Am J Physiol Heart Circ Physiol, 2017. 313(2): p. H265-H274.

17. Scott, I., et al., Identification of a molecular component of the mitochondrial acetyltransferase programme: a novel role for GCN5L1. Biochem J, 2012. 443(3): p. 655–61.

18. Thapa, D., et al., Loss of GCN5L1 in cardiac cells limits mitochondrial respiratory capacity under hyperglycemic conditions. Physiol Rep, 2019. 7(8): p. e14054.

19. Manning, J.R., et al., Loss of GCN5L1 in cardiac cells disrupts glucose metabolism and promotes cell death via reduced Akt/mTORC2 signaling. Biochem J, 2019. 476(12): p. 1713–1724.

20. Manning, J.R., et al., Cardiac-specific deletion of GCN5L1 restricts recovery from ischemia-reperfusion injury. J Mol Cell Cardiol, 2019. 129: p. 69–78.

21. Mori, J., et al., ANG II causes insulin resistance and induces cardiac metabolic switch and inefficiency: a critical role of PDK4. Am J Physiol Heart Circ Physiol, 2013. 304(8): p. H1103–13.

22. Chouchani, E.T., et al., Ischaemic accumulation of succinate controls reperfusion injury through mitochondrial ROS. Nature, 2014. 515(7527): p. 431–435.

23. Tao, R., et al., Sirt3-mediated deacetylation of evolutionarily conserved lysine 122 regulates MnSOD activity in response to stress. Mol Cell, 2010. 40(6): p. 893–904.

24. Hirschey, M.D., et al., SIRT3 deficiency and mitochondrial protein hyperacetylation accelerate the development of the metabolic syndrome. Mol Cell, 2011. 44(2): p. 177–90.

25. Kendrick, A.A., et al., Fatty liver is associated with reduced SIRT3 activity and mitochondrial protein hyperacetylation. Biochem J, 2011. 433(3): p. 505–14.

26. Liu, Y., G. Fiskum, and D. Schubert, Generation of reactive oxygen species by the mitochondrial electron transport chain. J Neurochem, 2002. 80(5): p. 780–7.

27. McLennan, H.R. and M. Degli Esposti, The contribution of mitochondrial respiratory complexes to the production of reactive oxygen species. J Bioenerg Biomembr, 2000. 32(2): p. 153–62.

28. Turrens, J.F. and A. Boveris, Generation of superoxide anion by the NADH dehydrogenase of bovine heart mitochondria. Biochem J, 1980. 191(2): p. 421–7.

29. Karamanlidis, G., et al., Mitochondrial complex I deficiency increases protein acetylation and accelerates heart failure. Cell Metab, 2013. 18(2): p. 239–50.

30. Vazquez, E.J., et al., Mitochondrial complex I defect and increased fatty acid oxidation enhance protein lysine acetylation in the diabetic heart. Cardiovasc Res, 2015. 107(4): p. 453–65.

31. Oberley, L.W. and T.D. Oberley, Role of antioxidant enzymes in cell immortalization and transformation. Mol Cell Biochem, 1988. 84(2): p. 147–53.

